# Decoding Speech and Music Stimuli from the Frequency Following Response

**DOI:** 10.1101/661066

**Authors:** Steven Losorelli, Blair Kaneshiro, Gabriella A. Musacchia, Nikolas H. Blevins, Matthew B. Fitzgerald

## Abstract

The ability to differentiate complex sounds is essential for communication. Here, we propose using a machine-learning approach, called classification, to objectively evaluate auditory perception. In this study, we recorded frequency following responses (FFRs) from 13 normal-hearing adult participants to six short music and speech stimuli sharing similar fundamental frequencies but varying in overall spectral and temporal characteristics. Each participant completed a perceptual identification test using the same stimuli. We used linear discriminant analysis to classify FFRs. Results showed statistically significant FFR classification accuracies using both the full response epoch in the time domain (72.3% accuracy, p < 0.001) as well as real and imaginary Fourier coefficients up to 1 kHz (74.6%, p < 0.001). We classified decomposed versions of the responses in order to examine which response features contributed to successful decoding. Classifier accuracies using Fourier magnitude and phase alone in the same frequency range were lower but still significant (58.2% and 41.3% respectively, p < 0.001). Classification of overlapping 20-msec subsets of the FFR in the time domain similarly produced reduced but significant accuracies (42.3%–62.8%, p < 0.001). Participants’ mean perceptual responses were most accurate (90.6%, p < 0.001). Confusion matrices from FFR classifications and perceptual responses were converted to distance matrices and visualized as dendrograms. FFR classifications and perceptual responses demonstrate similar patterns of confusion across the stimuli. Our results demonstrate that classification can differentiate auditory stimuli from FFR responses with high accuracy. Moreover, the reduced accuracies obtained when the FFR is decomposed in the time and frequency domains suggest that different response features contribute complementary information, similar to how the human auditory system is thought to rely on both timing and frequency information to accurately process sound. Taken together, these results suggest that FFR classification is a promising approach for objective assessment of auditory perception.

## 1 Introduction

Sound identification is a central feature of human communication. To successfully identify and assign appropriate labels to different sounds, individuals must be able to distinguish between them based on their spectro-temporal acoustic characteristics. Discrimination of different speakers and instruments is based on the perception of more complex spectro-temporal features called timbral qualities. These include the shape of the spectral envelope over time, the center frequency of sound spectrum, and frequency range emphasis (e.g., high versus low overtones). Finally, pitch perception in both speech and music most often varies proportionally with the fundamental frequency (F0). Given the importance of these acoustic characteristics for sound identification, it is thought that speech understanding therefore requires distinct and precise neural representations for different acoustic signals. In many instances, the integrity of these neural representations is inferred by tests of speech understanding or sound identification, such that good performance on these measures is generally thought to reflect neural encoding of sufficient integrity. Such tests are widely used in clinical audiologic practice when assessing performance of individuals with normal hearing and hearing loss (Lawson & Peterson, 2011). However, some individuals (e.g., young children) may not be able to reliably respond to signals, or—in the case of significant hearing loss—may have difficulty reporting what is heard with their hearing aid or cochlear implant. For these populations, an objective measure of the neural representation is of considerable interest for both clinicians and researchers.

In human listeners the neural representation of speech and non-speech sounds is often investigated using auditory evoked potentials (AEPs) (Atcherson & Stoody, 2012; Hall III, 2007c). In clinical audiologic practice, the auditory brainstem response (ABR) (Jewett et al., 1970; Hood et al., 1991; Hall III, 2007e; Brueggeman & Atcherson, 2012; Davis et al., 1985) and the auditory steady-state response (ASSR) (Galambos et al., 1981; Stapells et al., 1984; Hall III, 2007b; Strickland & Needleman, 2012) are widely used within an audiologic battery to detect the presence and degree of hearing loss. While these measures are excellent at helping to identify the degree and configuration of hearing loss, they provide little information regarding sound identification. For these reasons, discriminability of speech and non-speech sounds has been investigated through two specific components of cortical AEPs, called the mismatch negativity (MMN) and P300. Both of these potentials are elicited by an ‘oddball’ paradigm in which a sequence of standard sounds is interrupted by an infrequent change along any acoustic dimension (e.g., frequency, intensity, duration, location; for review, see Atcherson & White (2012)). The neural response to the infrequent stimulus differs from that observed to the frequent stimulus, and the size of this change has been shown to reflect the discriminability of these stimuli (Näätänen & Alho, 1995, 1997). MMN provides a neurophysiological measure of acoustic change detection when sounds are not attended to (Näätänen et al., 1993) and the P300 is modulated by attention to such changes (Davis (1964); for review, see Hall III (2007d)). The MMN is limited in use and scope because not all people exhibit this potential despite normal discriminability performance. In addition, both the MMN and P300 are modulated by state of arousal and diminished during sleep, making it difficult to test in infant populations.

One AEP candidate which may be better suited for determining whether the neural response of a signal is sufficient for identification on an individual level is the frequency following response (FFR, for review see N. A. Kraus et al. (2017)). The FFR is similar to the ABR in that both evoked potentials are recorded in the same manner, and generators that produce them overlap— namely the cochlea, VIIIth cranial nerve and inferior colliculi (Smith et al. (1975); Gardi et al. (1979); Sohmer et al. (1977); for review see Hall III (2007a)). The responses differ in that while the transient peaks of the ABR resolve after about 10 msec, the peaks of the FFR closely ‘follow’ the stimulating frequency in a steady-state response for as long as the stimulus continues (Hoormann et al.(1992); for review see Bhagat (2012)). The FFR has been used widely to investigate the neural encoding of complex tones (Greenberg et al., 1987), music (Bidelman & Krishnan, 2009; Musacchia et al., 2007), and speech sounds (Skoe & Kraus, 2010). Seminal studies into the FFR elicited by speech stimuli have shown that the strength of this response is greatest when auditory stimuli are correctly classified into speech categories (G. Galbraith et al., 1995) and is selectively enhanced to forward-running speech (G. C. Galbraith et al., 2004). Because the FFR is intimately related to the acoustics of the incoming signal, a number of potential applications have been suggested for its use. For example, the FFR is thought to be related to auditory experience, such that stronger FFRs are observed with greater degrees of musical skill (Musacchia et al., 2007; N. Kraus & Strait, 2015; Parbery-Clark et al., 2012; Wong et al., 2007) or language experience (Krishnan et al., 2005; S.Wang et al., 2016; Song et al., 2008). Similarly the FFR strength has been correlated with the ability to understand speech in the presence of background noise, and with development in infants (Anderson et al., 2015; Musacchia et al., 2018) and children (Russo-Ponsaran et al., 2004; Banai et al., 2007). Overall, these studies demonstrate the potential of the FFR to examine the integrity of neural coding of speech signals. However, most of these reports are correlational in nature (e.g., comparing FFR strength with some other variable) and do not address the question of sound identification.

One approach to predicting sound identification from AEP data is the use of classification. This refers to an analysis technique which attempts to predict a stimulus label or cognitive state from the corresponding brain response (Donchin, 1969; Blankertz et al., 2001). One advantage of classification is that it can be applied to responses to multiple stimuli, which more closely emulates what is required of participants when performing tests of speech identification. The classification approach also enables complex, high-dimensional data to be analyzed in its entirety, or in spatial and/or temporal subsets using a ‘searchlight’ approach (Su et al., 2012; Kaneshiro et al., 2015). To date, classification has been widely applied to cortical magnetoencephalography (MEG) and electroencephalography (EEG) data to study perception of visual object categories (Simanova et al., 2011; Carlson et al., 2013; Kaneshiro et al., 2015), and to a lesser extent music (Schaefer et al., 2011; Sankaran et al., 2018).

Classification-based approaches have also been applied to neurophysiological correlates of speech identification using FFR and AEP data. Results show that the spectral amplitude of the FFR in F0 and F1 bands can be used to correctly classify vowels with up to 70–80% accuracy (Sadeghian et al., 2015) and that spectral information related to the F2 band can be used to classify cortical evoked responses to vowels on the basis of single-trial data (Kim et al., 2014). Data acquired with an innovative combination of single-trial classification and machine-learning methods support the notion that the spectral amplitude of the FFR may be used to correctly predict vowel categorization into learned and novel vowel categories (Yi et al., 2017). Moreover, temporal information contained in the phase of theta oscillations (2–9 Hz) could correctly classify eight phonetic categories such that confusion matrices from phase and perceptual responses were not statistically distinguishable from one another (R. Wang et al., 2012). Finally, a recent report demonstrated that vectors of timing from cortical responses could be combined with vectors of spectral information derived from the FFR to successfully classify brain responses into speech categories in patients with mild cognitive impairment (Bidelman et al., 2017).

Taken together, these data support early phonetic classification research (Näätänen et al., 1978), and provide strong evidence that AEPs, and in particular the FFR, have potential to accurately predict speech identification when classification algorithms are applied. However, the components of the neural response underlying the classification results are poorly understood. For example, most investigations focused their classification analyses on spectral and temporal methods independently; only Lee & Bidelman (2017) appears to have combined both into their identification algorithm. Thus, it is unclear precisely which components of the FFR contribute to accurate classification of signals. Moreover, the range of stimuli which have been used to acquire FFR data for classification is rather narrow to date and has consisted largely of vowels. This is relevant because speech consists of both consonants and vowels and varies considerably in its spectral content over time (e.g., transitioning from consonants to vowels (CV) within a given phoneme). In such cases, it is unclear whether the accuracy of a classification approach changes over time, and if so, how the time course of those changes manifest. Finally, the extent to which classification can identify stimuli of completely different types is uncertain. For example, can responses to speech stimuli and musical notes be separated, and appropriate labels applied to each type? Here we addressed these questions by classifying FFR responses to CV phonemes and musical stimuli, and comparing the output of the classifier to the results of a perceptual-identification task. We then decomposed the FFR in order to identify which features of the neural response drive classification. Finally, we employed a temporal searchlight approach to classify these stimuli at different time points of the response with the intent of relating accuracy of these classification windows to acoustic features of the speech and music stimuli.

## 2 Methods

### 2.1 Participants and Stimuli

This research protocol was approved by Stanford University’s Institutional Review Board. All participants provided written informed consent before engaging in any research activities. Thirteen adult participants (7 female) participated in the study. Participants ranged in age from 20–35 years (mean 24 years), were fluent in English, and had no cognitive or decisional impairments. Normal hearing was confirmed with audiological hearing thresholds <20 dB HL across octave frequencies ranging from 500–4000 Hz. Each participant completed a general demographic and musical background questionnaire.

The stimulus set comprised three CV phonemes—*ba, da, di* —and three musical notes labeled *piano, bassoon*, and *tuba* based on their timbral qualities. Time- and frequency-domain visualizations of the stimuli are shown in Panel A of Figure 1. These stimuli were chosen due to their availability in the system used for data collection—advancing our long-term goal of clinical feasibility—as well as on the basis of their shared and distinct auditory features. For example, *ba* and *da* are identical in sustained vowel content but differ in their initial consonant, and can thus be distinguished perceptually on the basis of a formant transition. On the other hand, *da* and *di* share similar (though not acoustically identical) initial consonants and differ in the sustained vowel. The musical stimuli vary in their onset characteristics as well as the amplitude and phase structure of their harmonics. The F0 of each stimulus ranged from 97–107 Hz (*ba* 100.1 Hz, *da* 100.1 Hz, *di* 106.7 Hz, *piano* 98.3 Hz, *bassoon* 100.0 Hz, *tuba* 97.3 Hz), and remained consistent across the sustained portion of each waveform. We made minor modifications to the stimuli using Audacity^1^ software, first trimming each waveform to 135 msec—a reasonable duration for collection of FFR responses (Russo-Ponsaran et al., 2004; Bidelman & Krishnan, 2009; Coffey et al., 2016; Skoe & Kraus, 2010)—and then applying a linear fade out over the last cycle of each waveform to eliminate offset transients.

**Figure 1:**
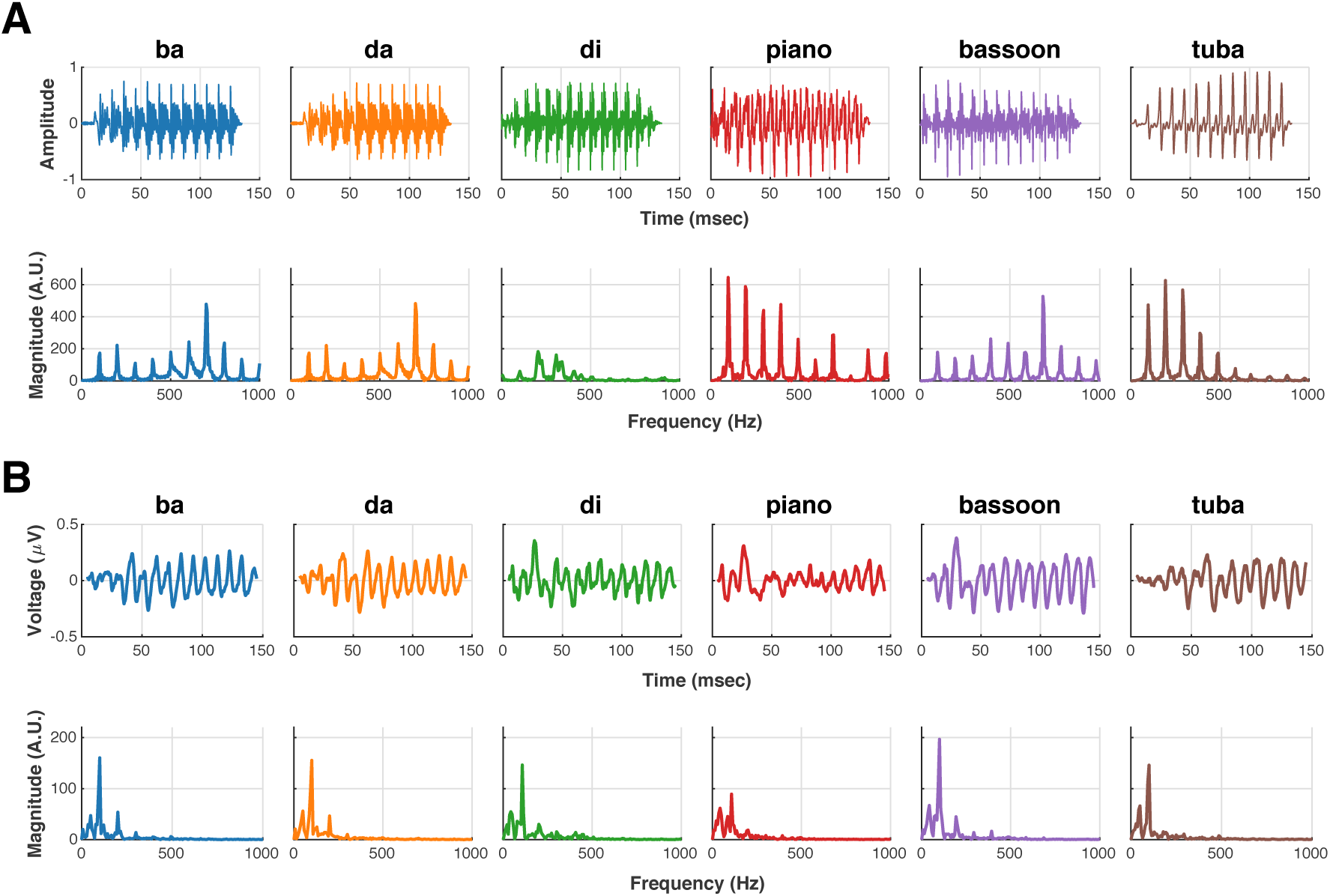
Stimulus and response visualizations. (A) Stimuli are 135 msec in length and include three consonant-vowel phonemes—*ba, da, di* —and three musical notes representative of different instruments— *piano, bassoon*, and *tuba*. Stimulus F0s are all near 100 Hz, but stimuli vary in fine temporal structure and harmonic content. (B) Global-average FFR responses (N=13; 2,500 responses per stimulus per participant). Responses vary in temporal and spectral composition; the strongest spectral peaks occur at F0 for every stimulus.

### 2.2 Data Collection

Each participant completed an FFR recording session and perceptual test. As even short-term training has been shown to modulate FFR responses (Wong et al., 2007; Song et al., 2008; Carcagno & Plack, 2011; Skoe et al., 2013; Russo et al., 2005) the FFR session was always conducted first. The combined sessions, including hearing screen and application of sensors, took approximately two hours for each participant.

The FFR session involved six recording blocks, with a single stimulus presented in each block. Ordering of blocks was randomized for each participant. Stimuli were presented with a 70-msec inter-stimulus interval (ISI, i.e., a silent interval of 70 msec between the conclusion of one stimulus and onset of the next) in alternating polarity. During recording, the participant was seated in a chair in a dimly lit room. A film without sound was presented to reduce participant movement during data acquisition; participants were also allowed to sleep during the recordings. Stimuli were played binaurally at 75 dB(C) via electrically shielded Etymotic ER-3 Insert Earphones.^2^

FFR responses were collected using the Advanced Research Module of the Intelligent Hearing Systems (IHS) SmartEP platform,^3^ which is approved for clinical use. Electrodes were placed at the frontal midline (Fz) with nasion reference and ground on bilateral earlobes. Electrode impedance was measured under 5 kΩ at the start of each recording session. FFR responses were recorded at a sampling rate of 20 kHz, band-pass filtered at acquisition between 30–1,500 Hz and segmented from -20.9–183.85 msec relative to stimulus onset. All FFRs were collected with an ongoing artifact rejection criterion of ±35 *µ*V. In each block, recording continued until 2,500 usable responses to stimulus repetitions—or *sweeps*—were obtained. A block was restarted if more than 250 rejections (10% of total sweeps) occurred.

As the goal of this study is to establish the feasibility of FFR classification under intact perception, we additionally included a perceptual test in order to verify that participants could behaviorally discriminate among the stimuli. The perceptual test began with a short familiarization phase designed to help participants associate a label with a given sound, and to familiarize them with the task. Here, the stimulus set was presented five times in order while the corresponding stimulus label (phoneme or instrument name) was shown onscreen. The participant then completed the test phase. In a test trial, a stimulus was played with no corresponding label shown, and the participant sub-sequently indicated the perceived label of that stimulus. Responses were given in a six-alternative forced-choice (6AFC) paradigm without feedback. Each stimulus was presented 20 times for a total of 120 test trials; trial ordering was randomized for each participant. The perceptual-identification task was written in Matlab^4^ using the Psychophysics Toolbox (Brainard, 1997) and was completed on a laptop; stimuli were played through Sennheiser HD 650 headphones.

### 2.3 FFR Data Export and Preprocessing

FFR data were exported from the IHS system on a per-participant, per-stimulus basis in 100-sweep averages—the minimum number of sweeps that our system could average for export. These FFRs were calculated by averaging an equal number of responses to a stimulus presented in alternating polarity, which Aiken & Picton (2008) define as the envelope FFR. Across all participants and stimuli, this produced 1,950 100-sweep averages (325 per stimulus). All subsequent preprocessing and analysis was performed in Matlab. We epoched the response data to a time interval between 5– 145 msec after stimulus onset to account for the latency of the FFR (Glaser et al., 1976; Hoormann et al., 1992)—resulting in a vector of 2,801 time samples—and then centered each epoched average via subtraction of the mean. We henceforth refer to these epoched, centered 100-sweep averages as *trials*.

All experimental data are made available under a CC-BY 3.0 License from the Stanford Digital Repository (Losorelli et al., 2019).^5^ Analysis code and preprocessed data are available on GitHub (also under a CC-BY 3.0 license).^6^ Organization of data and code files is detailed in Supplementary Figure S1, and mapping of participant identifiers from raw to preprocessed data files is documented in Supplementary Table S1.

### 2.4 FFR Classification

Classification is a machine-learning task which aims to assign correct categorical labels to observations of data. In the current context, a classifier is *trained* by building a statistical model from FFRs (observations) and their respective stimulus identifiers (labels). Then, in the *test* phase, FFR observations are input to the classifier without labels, and the classifier returns the predicted labels. We compare the labels predicted by the classifier with the actual labels of the test observations in order to compute the classification *accuracy*, which for this study is the percentage of test observations with correctly predicted test labels.

In order to determine how a classifier will perform in a predictive setting, it is good practice to exclude test observations from the training phase of building the model. One way to achieve this is by performing *cross validation*. In this procedure, the data are first divided into non-overlapping subsets—or *folds*—of roughly equal size. Classification is subsequently performed in an iterative fashion, withholding each fold once for testing and training on the remaining folds. *K* folds thus implies *K* train-test iterations, and this is referred to as *K*-fold cross validation.

A compact representation of classifier performance, in addition to classification accuracy, is the confusion matrix. For the present study, element (*i, j*) of a confusion matrix denotes the number of test observations actually belonging to category *i* that were predicted to belong to category *j* by the classifier. Values on the diagonal represent numbers of correct classifications. Our reported confusion matrices contain classifier predictions aggregated across all cross-validation folds.

We performed FFR classifications and visualized the results using the publicly available Mat-ClassRSA Matlab toolbox (B. C. Wang et al., 2017). As the classifier chose among 6 possible stimulus labels in this multi-category classification task, and we collected the same amount of response data to each stimulus, chance level was 1*/*6 (16.67%). All classifications used Linear Discriminant Analysis (LDA) (Hastie et al., 2009). To speed processing time of each classification, the dimensionality of the input data matrix was first reduced along the feature dimension (time samples or frequency bins) using Principal Components Analysis (PCA), retaining as many PCs were needed in order to explain 99% of the variance. In each cross-validation fold, PCA and number of PCs to retain was computed on only the training observations and then applied to the test observations. Statistical significance was assessed via permutation test (Golland & Fischl, 2003): Each classification was performed once on the intact data and 1,000 times with stimulus labels shuffled independently of the response data, and the corresponding *p*-value was computed by comparing the observed accuracy against the null distribution of permuted accuracies.

#### 2.4.1 Time-Domain Classification

For classification of FFRs in the time domain, the trial data from all participants were combined, and classification was performed using 10-fold cross validation. The trial-by-time matrix input to the classifier was of size 1,950×2,801, and the *features* describing each observation were time samples of response data. Trial ordering was randomized prior to partitioning for cross-validation in order to better distribute the participants’ data among the folds. In a follow-up classification, we further averaged the trials within-participant into *pseudo-trials* (Guggenmos et al., 2018) comprising 5 trials (500 sweeps) of response to a given stimulus, as used in Sadeghian et al. (2015). This produced an aggregated pseudo-trial-by-time matrix of size 390×2,801 across participants—leaving the feature dimension unchanged while decreasing the number of observations for classification (but likely improving their SNR). As classification of 500-sweep pseudo-trials was found to produce higher accuracy than classification of 100-sweep trials, we performed all subsequent classifications on 500-sweep representations of the data.

We next classified the data using a leave-one-participant-out (LOO) cross-validation scheme. Here, we performed 13-fold cross validation, where in each fold all observations from a single participant were withheld for testing, while the model was trained on the data from the remaining participants. We performed two such classifications. First, training and testing was performed using the within-participant pseudo-trials computed previously; then, pseudo-trials for training in each fold were computed across-participant on a per-stimulus basis, and the resulting model was tested on the within-participant pseudo-trials of the holdout test participant.

#### 2.4.2 Frequency-Domain Classification

One way in which the FFR differs from event-related cortical EEG responses is in its direct encoding of the auditory stimulus. Therefore, we speculated that FFR classification might be feasible in the frequency domain. To prepare the data for these analyses, we first computed the FFT of within-participant pseudo-trials. Next, taking into consideration the filter settings at data acquisition and the frequency range in which meaningful information is encoded by the FFR (Skoe & Kraus, 2010), we retained for further analysis only the 141 complex values corresponding to frequencies between 0–1,000 Hz.

A waveform can be fully characterized in the frequency domain via magnitude and phase values— or equivalently by its real and imaginary coefficients—at each frequency bin. For our first frequency-domain analysis, we classified this complex representation to confirm that accuracy would be comparable to time-domain classification accuracy. Here we used 10-fold cross validation, and the feature vector comprised real and imaginary Fourier coefficients (which occupy a shared data scale while phase and magnitude values do not) from 0–1,000 Hz, for a pseudo-trial-by-coefficient input matrix of size 390×282.

We next performed classifications on magnitude and phase alone. Fourier magnitude denotes the amount of energy at each frequency bin while phase describes precise timing information at each frequency. Therefore, if our previous classifications of complete responses were successful, these analyses could potentially elucidate the relative contributions of magnitude and phase in successful decoding. Over the same range of frequencies used above, we classified Fourier magnitudes, computed as the absolute values of each complex coefficient. We separately classified phase, computed as the angle at each frequency bin. Each of these classifications involved 10-fold cross validation and operated on an input pseudo-trial-by-feature (Fourier magnitude or phase) matrix of size 390×141.

#### 2.4.3 Temporal Searchlight Classification

Our analyses so far have considered time- and frequency-domain representations of the FFR over the full response epoch. With Fourier magnitude and phase classifications, we decomposed the FFR in order to determine whether classification accuracy could be explained by specific response features. For our final classifications, we decomposed the response in a different fashion, now over temporal subsets of the response epoch. This technique is referred to as temporal ‘searchlight’ (Su et al., 2012) and has been found to reveal useful information about the dynamics of visual object category processing in cortical EEG classification (Kaneshiro et al., 2015). For our searchlight analysis, we performed separate classifications on overlapping temporal windows 20 msec in length with a 10-msec (50%) overlap between windows, for a total of 13 windows. In each temporal window, we classified four representations of the response: Time domain (390×400 input matrix); frequency domain, real and imaginary coefficients (390×40 input matrix); frequency domain, magnitude only (390×20 input matrix); and frequency domain, phase only (390×20 input matrix). All classifications used 10-fold cross validation and the within-participant 500-sweep pseudo-trials used previously. We performed multiple comparison correction across the resulting 13 *p*-values using False Discovery Rate (Benjamini & Yekutieli, 2001).

### 2.5 Analysis of Perceptual Responses

While FFR confusion matrices were constructed by means of a computational classification procedure, in the perceptual case the classification is effectively performed by each participant in their reporting of perceived stimulus categories. Therefore, the perceptual confusion matrices could be constructed by directly comparing each participant’s reported results with the actual stimulus labels of the trials. Reported global perceptual accuracy was obtained by aggregating confusion matrices across participants. We assessed statistical significance of perceptual responses by comparing observed results with a null distribution of accuracies obtained by randomly permuting actual labels 1,000 times for each participant.

### 2.6 Visualization of Confusion Matrices

As the confusion matrix provides a more comprehensive summary of a classifier’s performance than accuracy alone, we visualized the confusion matrix for every classification. In addition, classifier confusions can be treated as measures of similarity among the classes (Shepard, 1974), and consequently a confusion matrix can be treated as a proximity matrix, converted to a distance matrix, and visualized hierarchically as a dendrogram. We created dendrograms from confusion matrices using the procedure outlined in (Kaneshiro et al., 2015): The distance matrix was created by first scaling each row of the confusion matrix by its respective diagonal entry (achieving unity self-similarity), symmetrizing the matrix using the geometric mean, and finally computing distances linearly as 1*-*similarity; following this, hierarchical clustering was performed using unweighted pair grouping method with averaging (UPGMA) linkage. In the resulting tree visualizations, the y-axes denotes distance, and the height a given tree must be traversed in order to travel between any two classes is the distance between those classes.

## 3 Results

### 3.1 Time-Domain Classification

For our first analyses, we classified FFRs in the time domain. To start, we input what was our closest representation of ‘single-trial’ FFRs—the 100-sweep averaged trials output by the IHS system—and performed 10-fold cross validation. Here, the mean accuracy across cross-validation folds was 61.4% (*p* < 0.001) compared to chance level of 16.67%. The next classification of 500-sweep pseudo-trials was more successful, with a mean accuracy of 72.3% (*p* < 0.001). The confusion matrix and dendrogram of the pseudo-trial classification are shown in Panel A of Figure 2. Examination of the confusion matrix indicates that not all stimuli were decoded with equal accuracy. Rather, the classifier tended to confuse responses to *ba* and *da*, as shown by large values in the off-diagonal. Responses to *di* and *piano* also tended to be confused, though to a lesser extent, while responses to *bassoon* and *tuba* classified best (classwise accuracies of 87.7% and 90.8%, respectively). The accompanying dendrogram makes the structure of the confusion-based similarities more clear, with *ba*/*da* forming the tightest category cluster (smallest distance), followed by *di* /*piano*.

**Figure 2:**
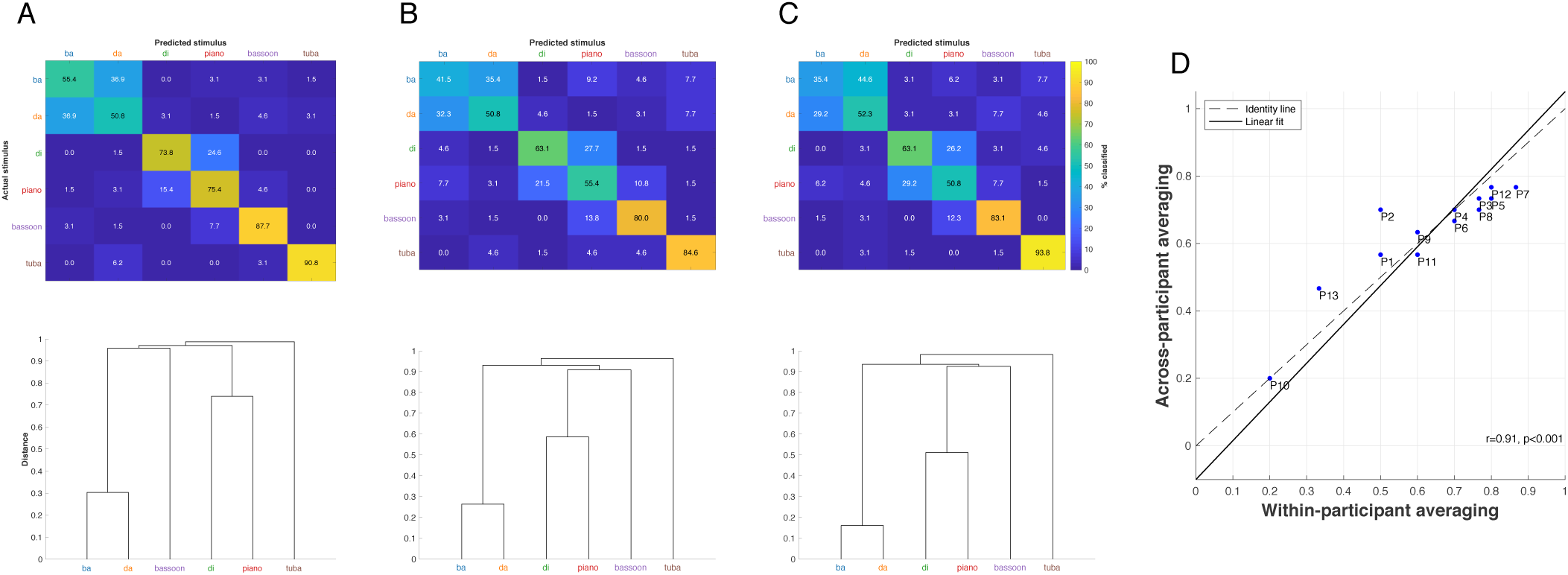
Time-domain classification results. Classification results for 500-sweep pseudo-trials, visualized as confusion matrices and dendrograms. (A) Pseudo-trials were computed within-participant, and across-participant classification was performed using 10-fold cross validation. Mean accuracy was 72.3% (*p* <0.001). Greatest confusion occurred between responses to *ba* and *da. Tuba* responses classified with highest accuracy. (B) Single participants were withheld for testing during (13-fold) cross validation. Pseudo-trials of the training data were computed within-participant. Mean accuracy was 62.6% (*p* <0.001). (C) Single participants were withheld for testing during (13-fold) cross validation. Pseudo-trials of the training data were computed across-participant. Mean accuracy was 63.1% (*p* <0.001). (D) Scatter plot of within-participant versus across-participant accuracies for single test participant accuracies from (B) and (C).

In a clinical setting, the predictive power of classification becomes especially relevant for assessing responses from previously unseen patients. To explore the feasibility of this scenario, we next iteratively trained the classifier on data from all but one participant and then tested on the data from that holdout participant. As participant-specific attributes of the test data cannot be taken into account during training, this is a more challenging task. However, it also more closely resembles the application of FFR classification in a real-world setting. We performed these classifications in two ways, first training the model on pseudo-trials averaged within-stimulus and within-participant, and next on pseudo-trials averaged within-stimulus but across-participant. When pseudo-trials of the training partitions were averaged within-participant (Figure 2B), mean classifier accuracy was 62.6% (*p* <0.001); results were similar when training pseudo-trials were computed across-participant, with an overall classification accuracy of 63.1% (*p* <0.001) (Figure 2C). In both cases, the structure of the confusions as shown in the confusion matrices and dendrograms corresponded to that obtained when data from all participants were distributed among the training and testing folds (Figure 2A). Finally, a comparison of holdout participant accuracies between the two pseudo-trial averaging procedures (Figure 2D) indicated that accuracies from across-participant averaging were highly correlated with those from within-participant averaging (*rho* =0.91, *p* <0.001). Individual participant accuracies were statistically significant (*p* <0.05, FDR corrected) for all participants except P10.

### 3.2 Frequency-Domain Classification

While cortical EEG classification studies typically operate in the time domain of the response, for the present study we also explored whether FFRs could be classified in the frequency domain. For our first frequency-domain analysis, we input real and imaginary Fourier coefficients between 0–1,000 Hz to the classifier. As expected, the resulting classification accuracy was similar to that obtained with the time-domain response (74.6%, *p* <0.001). As can be seen in the confusion matrix and dendrogram in Panel A of Figure 3, the structure of similarities was also similar to that of the time-domain classification, with strongest similarity between *ba*/*da*, followed by *di* /*piano*.

**Figure 3:**
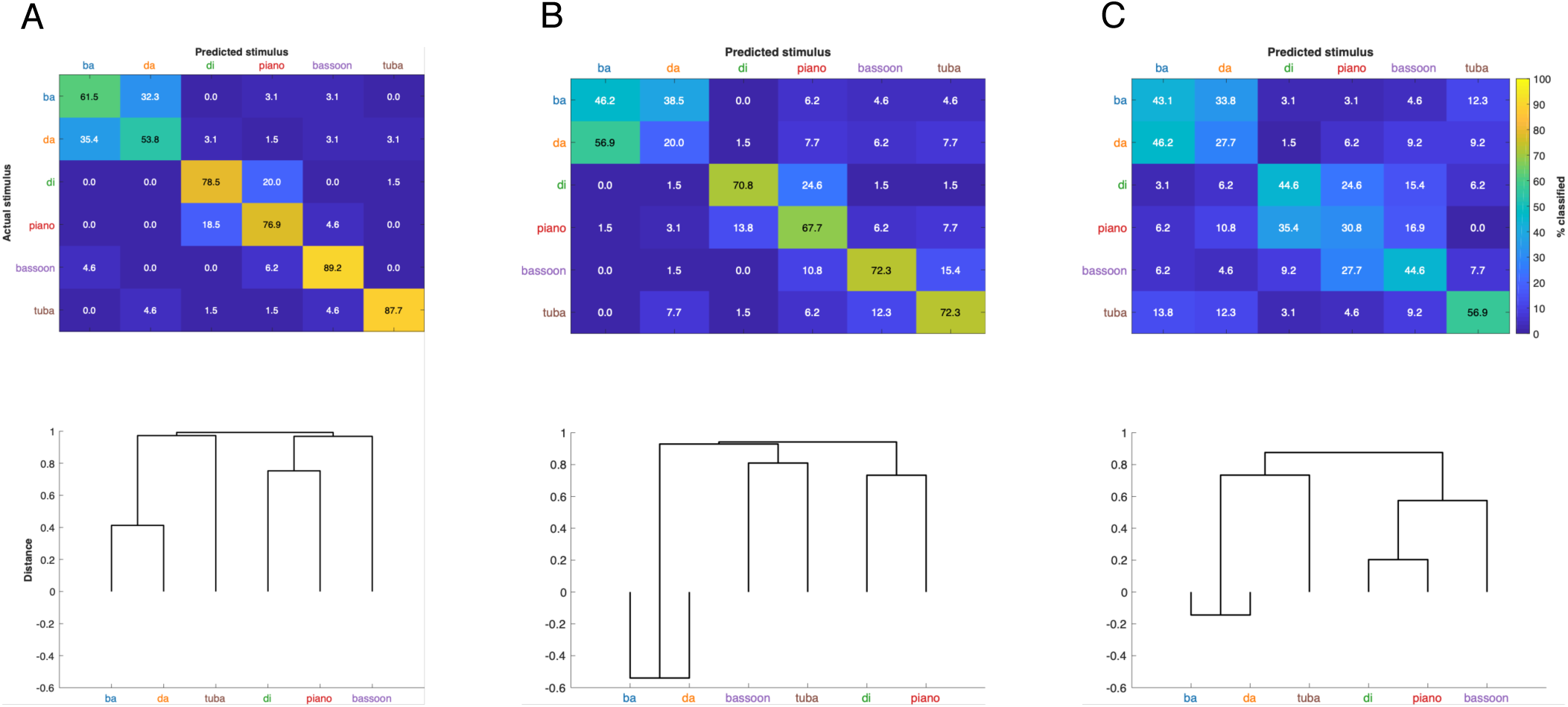
Frequency-domain classification results. Classification of 500-sweep pseudo-trials, computed within-participant, was performed using 10-fold cross validation and visualized as confusion matrices and dendrograms. (A) Classification of real and imaginary FFT coefficients up to 1,000 Hz. Mean accuracy was 74.6% (*p* <0.001). (B) Classification of FFT magnitudes up to 1,000 Hz. Mean accuracy was 58.2% (*p* <0.001). (C) Classification of FFT phase angles up to 1,000 Hz. Mean accuracy was 41.3% (*p* <0.001). For all feature representations, *ba* and *da* responses were confused most by the classifier.

We next decomposed the frequency-domain representation of the responses into Fourier magnitudes and phases for frequencies up to 1,000 Hz, and classified each of these representations separately. Classification of magnitudes produced an overall accuracy of 58.2% (*p* <0.001), while classification of phase values was less successful, with an overall accuracy of 41.3% (*p* <0.001). As is shown in the confusion matrix and dendrogram in Panels B and C of Figure 3, confusions between *ba* and *da* were exacerbated, with these two categories now displaying a negative distance (resulting from a greater number of misclassifications than correct classifications between the categories), while *di* /*piano* formed the second-closest category cluster and *tuba* classified most successfully in both cases. We note that the structure of the phase-classification dendrogram matches those of the time-domain and complex frequency-domain dendrograms, while classification of magnitudes produced a slightly different similarity structure.

### 3.3 Temporal Searchlight

Our final classifications were conducted on 20-msec temporal subsets of the response, with 10-msec overlap between time windows. This approach would highlight whether a particular range of time in the response—corresponding to the formant transition or to the sustained portion, for example— was especially essential to the success of classifications conducted across all time. Each temporal classification was performed in the time domain, on real and imaginary Fourier coefficients, and on Fourier magnitude and phase separately. Temporally resolved per-class and overall accuracies for time-domain classifications are shown in Figure 4A. Here, we observed a gradual decline in overall classification accuracy over the time course of the response. We also found that the stimuli varied in their decodability over time. For instance, *ba* and *da* were best decoded in the early portion of the response and decline thereafter, while *tuba* responses classified worst in the early response, but from the fourth time window (35–55 msec after stimulus onset) were the best-classifying responses among the stimulus categories.

**Figure 4:**
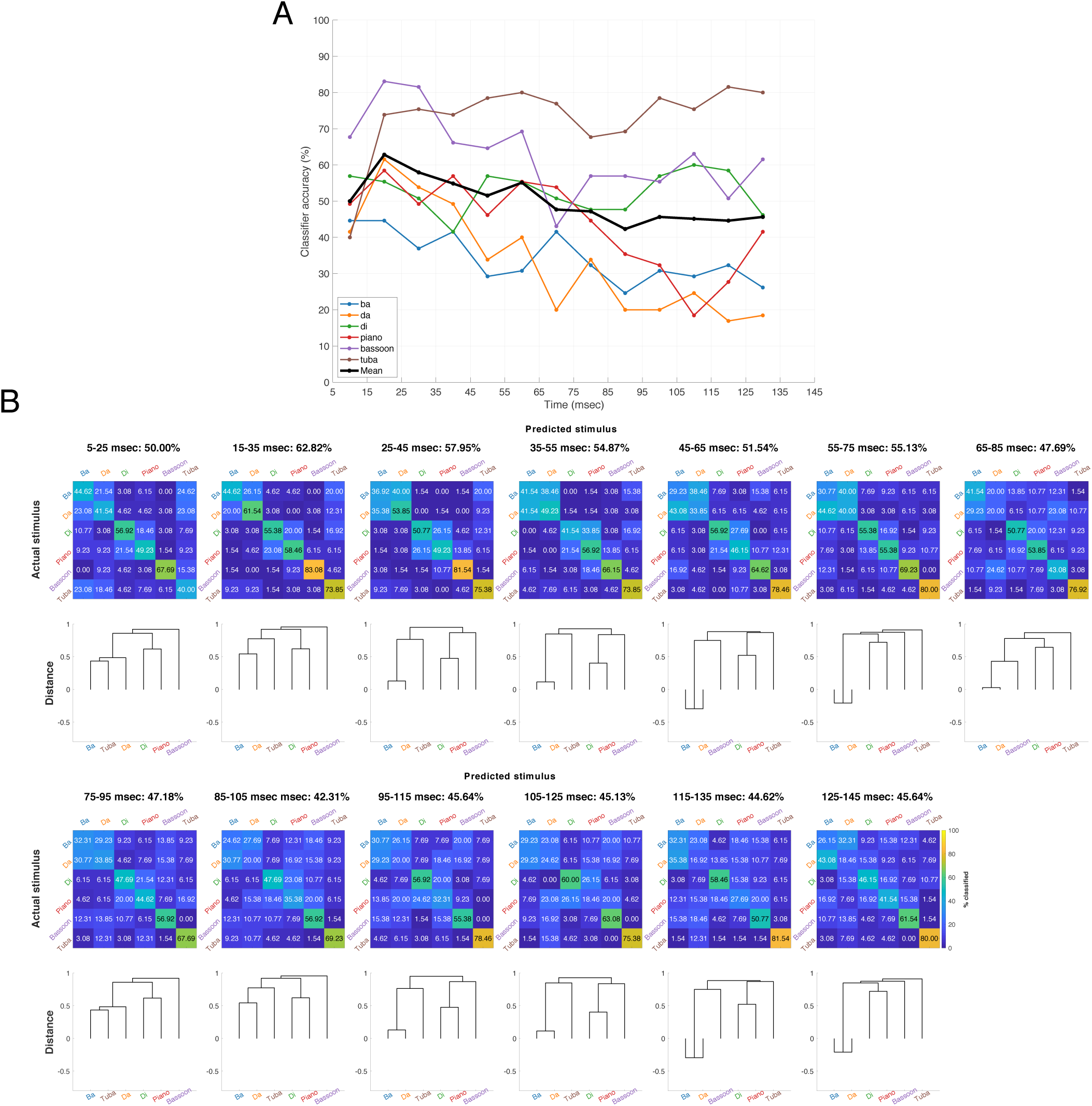
Temporal searchlight classification. Classifications were performed on temporal subsets of the time-domain response (500-sweep pseudo-trials, calculated within-participant), using 20-msec windows advancing in 10 msec increments. (A) Classifier accuracies of single categories over time with mean accuracies overlaid. (B) Mean accuracies, confusion matrices, and dendrograms for each temporal searchlight classification.

The accompanying confusion matrices and dendrograms provide insight into the stimulus structure over the time course of the neural response. Responses to *ba* and *da*, which have formed the closest category cluster in all analyses thus far, similarly clustered together here, with the notable exception of the first time window (5–25 msec after stimulus onset). Our secondary cluster of *di* /*piano* was generally present, although these two categories separated during a later section of the response.

As with classification of the full 5–145-msec window, temporal searchlight results for the real and imaginary coefficients up to 1,000 Hz performed similarly to the time-domain searchlight classifications. Temporal searchlight of Fourier magnitude and Fourier phase classifications performed worse than the fully characterized time-domain and frequency-domain responses. The results for the three frequency-domain searchlight classifications are included in Supplementary Figure S2–S4.

### 3.4 Perceptual Results

Each FFR recording was accompanied by a separate perceptual identification test. Given that intact stimuli were used in the experiment and all participants had normal hearing, we expected perceptual accuracy to be near 100%. Indeed, perceptual performance was high, with mean accuracy of 90.6% (*p* <0.001). As overall perceptual accuracies were high for each stimulus (84.9%–98.1%, Figure 5), we observed greater distance among all categories.

**Figure 5:**
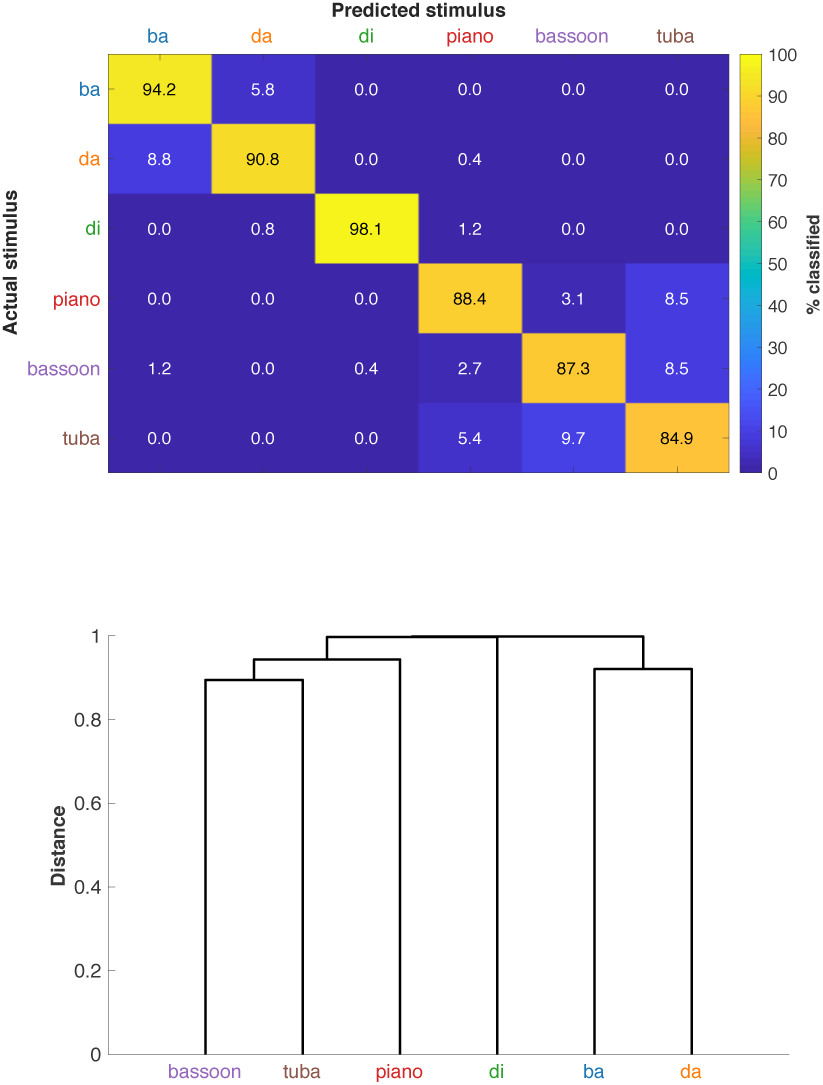
Perceptual identification results. Participants completed a 6AFC perceptual identification task separately from the FFR session. Mean confusion matrix and dendrogram across participants (N=13). Overall accuracy was 90.6% (*p* <0.001).

Perceptual accuracy of all but one participant exceeded 75% (Supplementary Figure S5). We did not observe a significant correlation between perceptual performance and leave-one-participant-out FFR classification performance when pseudo-trials were computed within-participant (*rho* =0.04, *p* =0.66).

## 4 Discussion

In this study we have demonstrated that FFRs elicited by both speech and music stimuli can be successfully classified, and that the pattern of classification approximates that observed with a perceptual-identification task. Here, overall accuracy on the perceptual-identification task was 90.6%, while the overall classifier accuracy was 72.3%. Our classifier accuracy for these CV phonemes and musical instruments is similar to that observed with vowels alone (∼70–80%; (Sadeghian et al., 2015), and contributes to a small but growing body of research on FFR decoding and novel analysis approaches.

Most FFR research to date has focused on descriptive approaches, which include obtaining the neural response by averaging thousands of stimulus presentations, and then comparing the FFR strength to performance on another task (see N. A. Kraus et al. (2017) for a review). In contrast, classification allows one to determine the extent to which an individual’s FFR responses can be correctly assigned stimulus category labels. This analysis approach thus more closely emulates the process of sound identification that humans perform repeatedly across the lifespan.

Our results suggest that overall classifier performance is heavily driven by accurate labeling of responses to musical instrument and *di* stimuli. For these stimuli, time-domain classwise accuracy ranged from 74% for *di* to 91% for *tuba* (Figure 2A). In contrast, responses to *ba* and *da* phonemes classified at 55.4% and 50.8%, respectively. While these accuracies exceed the six-class chance level of 16.7%, they are notably lower, and the majority of misclassifications occur between the two categories. One plausible explanation is that the difference in classifier accuracy for these FFRs reflects how robustly the acoustic characteristics of the signal are represented in the neural response. For example, *da* and *di* differ in the vowel portion of the CV phoneme, while *ba* and *da* differ largely in the formant transition from the consonant to the vowel but are identical in the sustained vowel region. Notably, the vowel-evoked FFR amplitude is greater and sustained over a longer time range than that of consonants (Skoe et al., 2015), which may contribute to a classifier that is more heavily weighted towards the vowel-evoked response. Future research could therefore study in greater detail the contribution of FFR encoding of steady-state and transient acoustic features to successful classification when these attributes are varied in a parametric fashion, in order to better understand the contributions of each.

We used different decompositions of our response in order to determine the contributions of the transient and steady state acoustics. First, we applied the classifier to 20-msec moving windows to assess temporal dynamics of FFR decoding across time. Results from these analyses suggest that accuracy is highest when the entire response is analyzed, regardless of which time frame was analyzed. However, relative accuracies of the temporal windows provide insight into the time-varying features of the stimulus that may be driving the classifier. Responses to *tuba*, for example, classify much better during the sustained portion of the response relative to the onset. In contrast, responses to *ba* and *da* are most successfully decoded in the region of the formant transition. These findings support the idea that similarities among stimuli are reflected in similarities of the neural response. Our conclusion is that for classification to optimally identify stimuli with more transient acoustic characteristics such as consonants, the feature vector passed into the classifier may need to be modified to emphasize specific components of the neural response more heavily (e.g., give higher frequencies heavier weighting in an effort to better capture transient changes associated with formant transitions).

To further understand the classification process used here, we not only decomposed the response in time, but also assessed classification accuracy according to frequency-domain features. We independently analyzed complex frequency values as well as phase and magnitude components of response spectra. Independent classification accuracy of Fourier magnitude and phase was 58.2% and 41.3%, respectively, compared to 72.3% accuracy of the global response (e.g., combined phase and magnitude). Our data suggest that the integration of magnitude and phase information, as well as the corresponding temporal characteristics, contributes to optimal classification accuracy with our method.

We have shown that classification can produce significant and interpretable results using at least one-fourth fewer stimulus repetitions than are required for averaging-based analyses (4,000–6,000 averages per trial; Skoe & Kraus (2010)). As multivariate approaches such as decoding can make use of multiple response features at once during analysis, they exhibit increased sensitivity compared to univariate averaging-based approaches (Haynes & Rees, 2006). The high accuracies obtained in the present study suggest that the classifier is discovering patterns in the observations that may not be readily distinguishable to human observers. This reduction in number of trials aligns with the approach taken in cortical M/EEG classification studies, many of which operate on single trials of response (Kaneshiro et al., 2015; Sankaran et al., 2018) as opposed to up to hundreds of trials needed for averaged ERP analyses (Woodman, 2010). In addition to classifying FFRs with 10-fold cross validation—in which each participant’s response data was distributed among training and test folds—we also carried out cross validation with single-participant test partitions in order to determine whether data from a participant who was not used to train the model could be successfully classified. Compared to the 10-fold case, results in this scenario showed an approximately 10% reduction in overall accuracy, for both within- and across-participant averaging of training observations; these results suggest that individual differences play a role in FFR classification.

The final observation is that, while the classification accuracy was clearly above chance, it still remained below that observed on the perceptual identification task. Both Sadeghian et al. (2015) and our results show classification for vowels occurring on approximately 70-80% of trials, as opposed to the high perceptual accuracy observed here for our CV stimuli (91-98%). This suggests that further refinement of the feature input to the classifier or classifier algorithm choice (i.e., how the classifer weighs particular features of the input) is necessary in order to identify speech signals at a rate of accuracy exhibited by that of individuals with normal hearing. Resolving this issue is a crucial step toward future clinical applications. In contrast to the speech signals, classifier and behavioral accuracy were more similar for the musical instruments (84.6% versus 86.9%, respectively). We speculate that the increased classifier accuracy in the music condition reflects the ability of the FFR to encapsulate spectro-temporal features needed to distinguish between instruments. The relationship between perceptual and FFR classifiers may also be impacted by the relative ability of listeners to correctly identify different musical instruments. Here, participants 10, 12, and 13 had perceptual-identification scores for the musical instruments which were much lower than those of the other participants (individual participant confusion matrices are shown in Supplementary Figure S5). Given that all participants had normal hearing, a plausible interpretation of these data is that the individuals could discriminate among the stimuli, but had inconsistent label mappings for the instruments. We conjecture that these results may not reflect the true perceptual abilities of the participant, but rather difficulties with assigning the correct instrument label.

We identify several opportunities for improvement in future research. First, and perhaps most important would be improvements to the classification algorithm itself. For example, improvements in feature selection or other aspects of the classifier could reduce the necessary data size for optimal classification, the optimal number of sweeps to be averaged in trials or pseudo-trials, or the total analysis time. Such improvements to the classifier could better account for individual differences when training the model. In the present study we did not have enough data to both train and test on individual participants. Reducing the necessary amount of data to build a reliable model could enable this step and is likely to prove crucial for modeling data from listeners with hearing loss who use hearing aids or cochlear implants. A second opportunity for improvement would be explore the extent to which small differences in stimuli influence the classification accuracy. Here, the stimuli differed slightly in F0 (97–107 Hz), raising the possibility that these small differences may have influenced the accuracy of our classifier. While we cannot rule out this possibility, we speculate that any effect was minimal because our classifier accuracy was largely similar to that observed with data obtained with F0-aligned stimuli (Sadeghian et al., 2015). Nonetheless, future research could examine FFR classifications for stimuli whose F0 frequencies are exactly matched, as in that case classification would be driven entirely by differences in spectral and temporal structure.

In closing, this study provides evidence that FFR classification can be employed to discriminate among responses to CV phonemes and musical instruments. In the present study, we verified that FFR classification is successful under intact perception, and explored the potential stimulus features that drive FFR decoding accuracy. These data add further evidence to a growing body of literature that EEG decoding approaches hold considerable promise for clinical applications in patients with hearing loss, or investigations of the developing auditory system when objective measures of behavioral abilities are desired.

## 5 Acknowledgments

The authors gratefully acknowledge Karanvir Singh and Vivian Lou for their assistance with data collection; Bernard Wang for adding requested features to the classification code; Duc Nguyen for her help in preparing the code for the perceptual identification test; and Malcolm Slaney for helpful feedback on a draft of the manuscript.

## Footnotes

1 http://www.audacityteam.org

2 https://www.etymotic.com/

3 https://www.ihsys.com/

4 https://www.mathworks.com/

5 https://purl.stanford.edu/cp051gh0103

6 https://github.com/slosorelli/FFR-classification-2019

