## Supplemental Figures for "Decoding Speech and Music Stimuli from the Frequency Following Response"

### Supplementary Materials

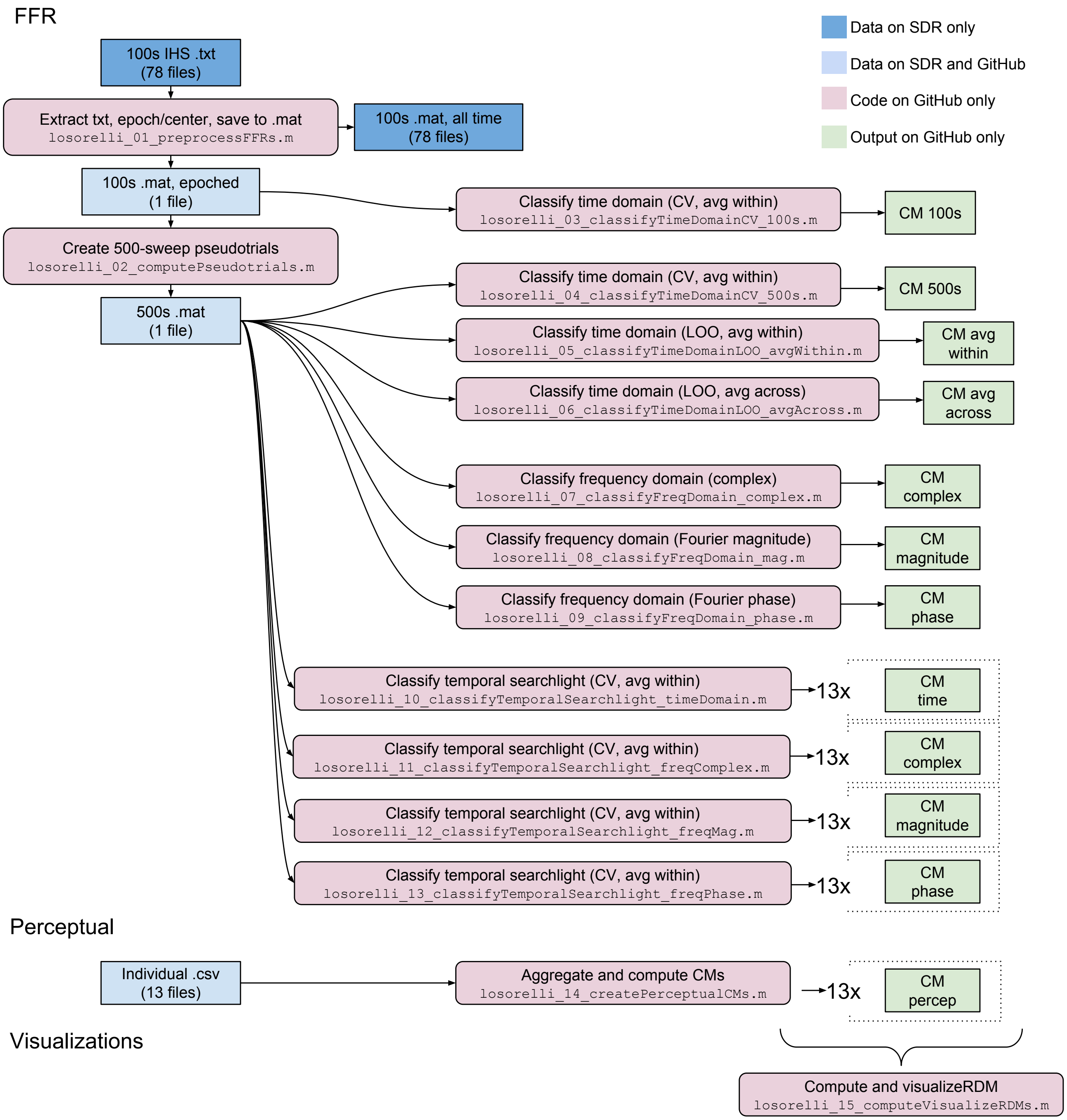


Figure S1: Organization of publicly available experimental data and analysis code.

| Analysis identifier | Raw identifier |
|---|---|
| P1 | S01 |
| P2 | S02 |
| P3 | S03 |
| P4 | S04 |
| P5 | S05 |
| P6 | S06 |
| P7 | S08 |
| P8 | S09 |
| P9 | S11 |
| P10 | S12 |
| P11 | S13 |
| P12 | S14 |
| P13 | S16 |

Table S1: Mapping of participant identifiers of preprocessed data (used for analysis) to those specified in raw .txt files of the IHS export.

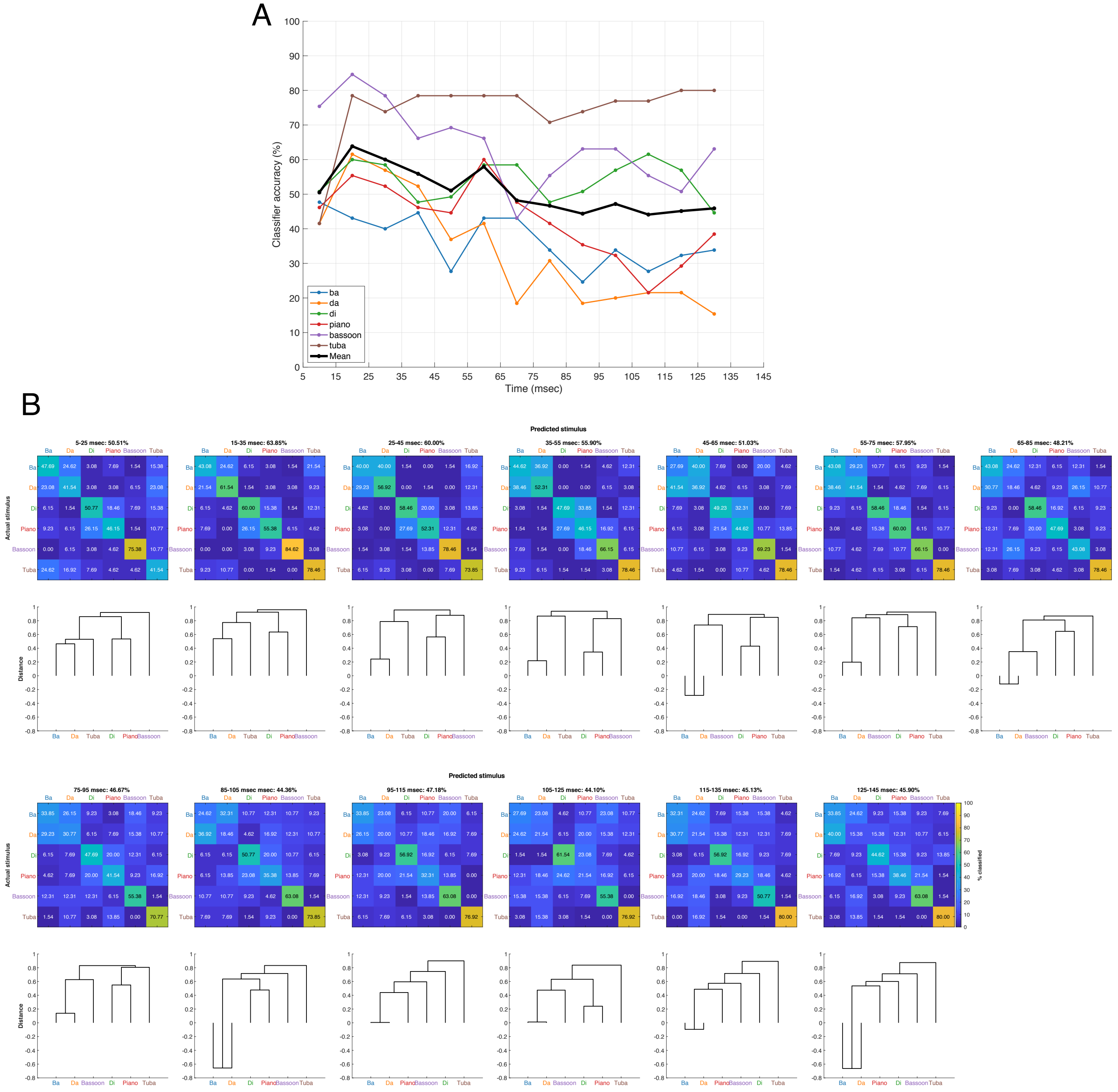


Figure S2: Temporal searchlight results for classification of real and imaginary Fourier coefficients.

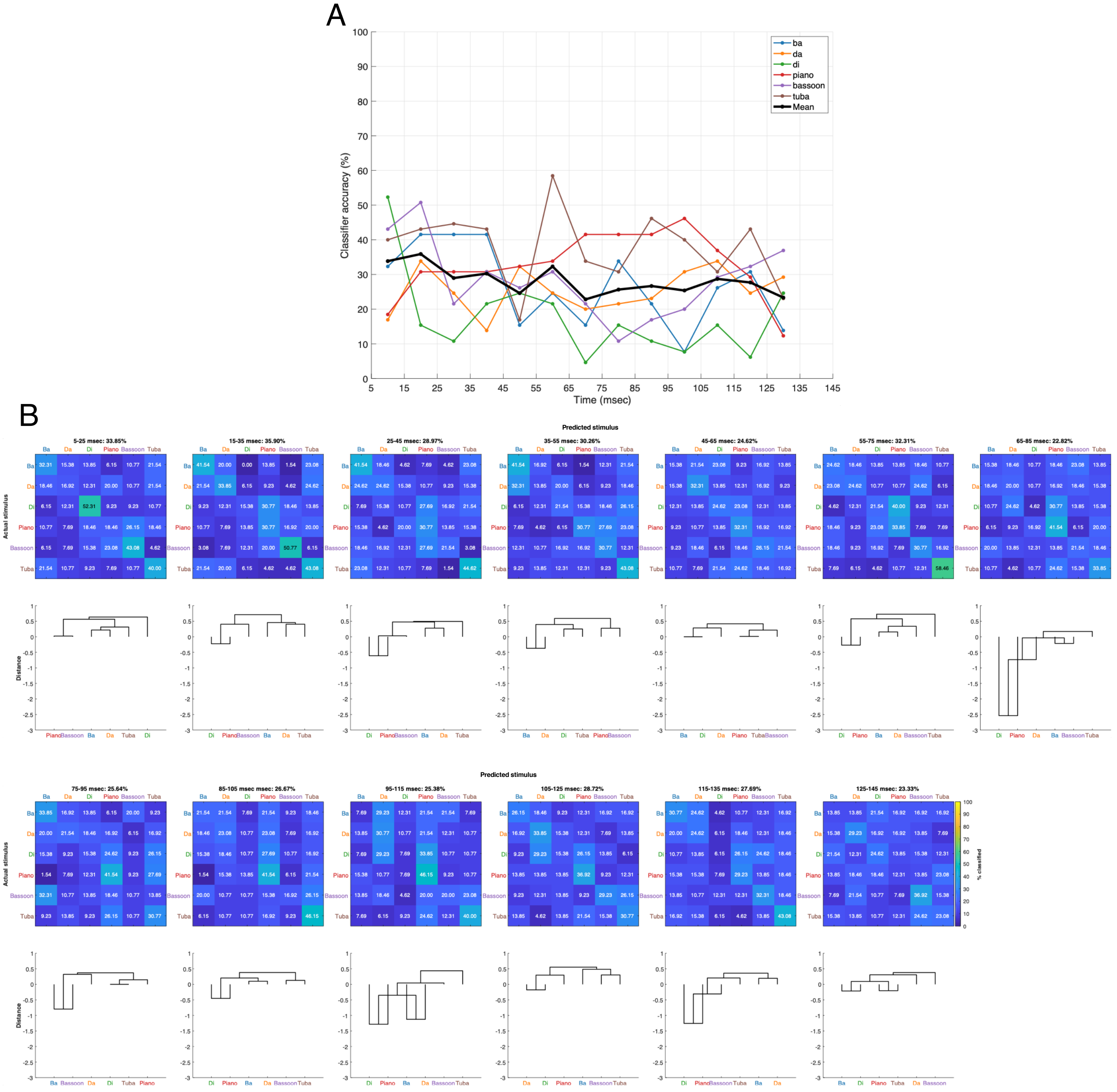


Figure S3: Temporal searchlight results for classification of Fourier magnitude.

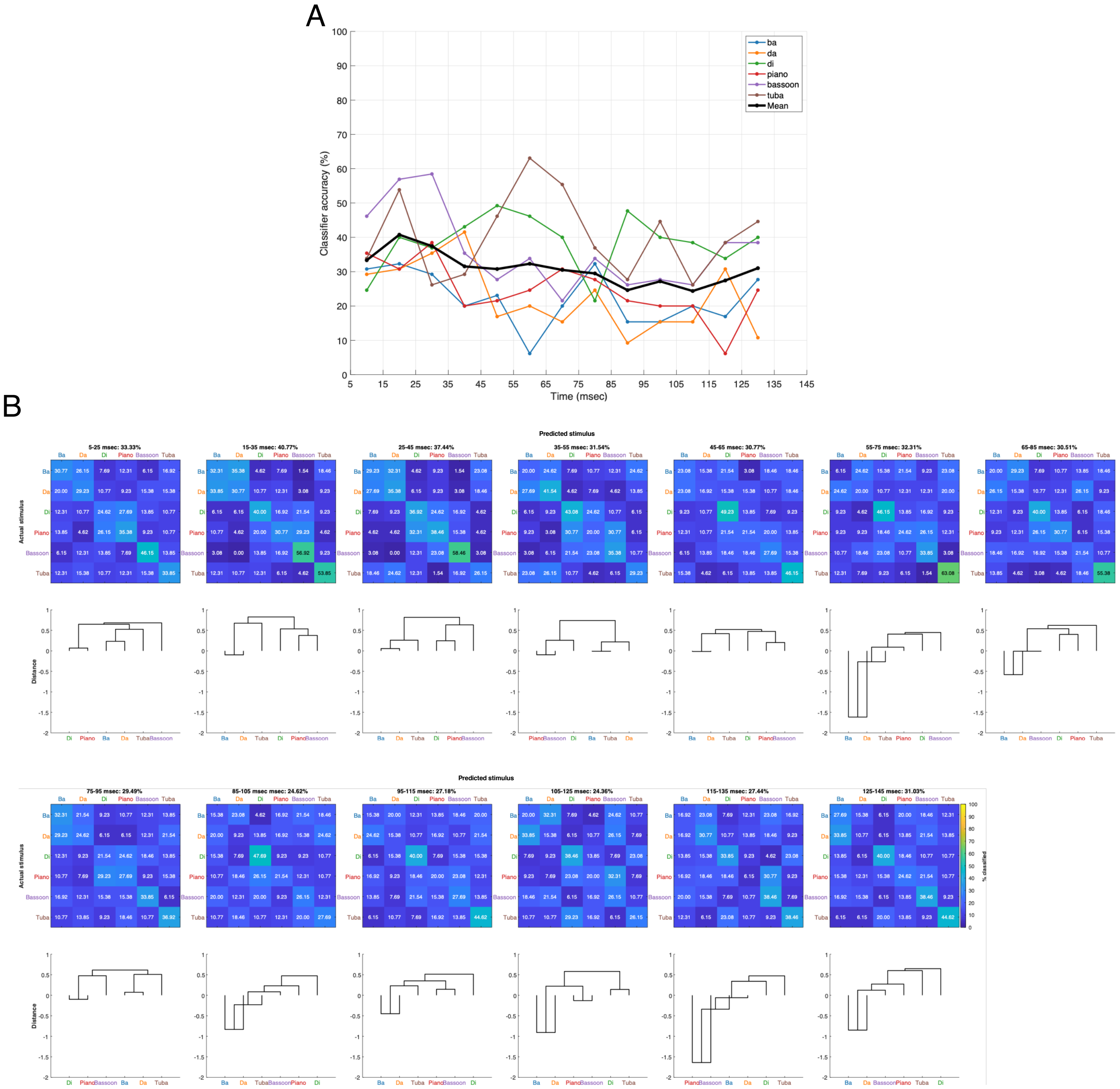


Figure S4: Temporal searchlight results for classification of Fourier phase.

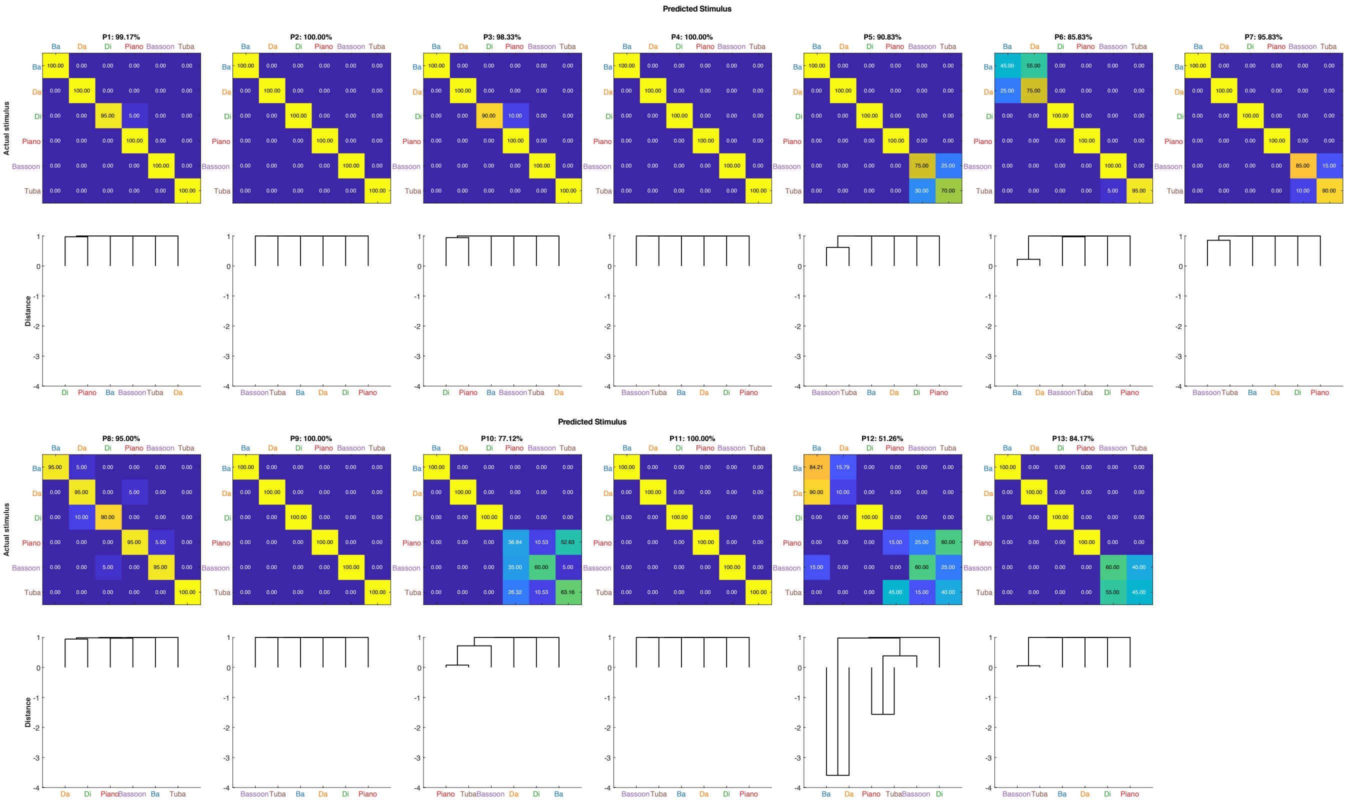


Figure S5: Perceptual confusion matrices from individual participants.
